# Increased Callosal Thickness in Early Trained Opera Singers

**DOI:** 10.1101/2025.02.28.640273

**Authors:** B. Kleber, C. Dale, A.M. Zamorano, M. Lotze, E. Luders, F. Kurth

## Abstract

Extensive research has shown how the corpus callosum adapts to early sensorimotor training in instrumental musicians, yet less is known about these effects in professional singers. This study used high-resolution MRI to investigate variations in callosal thickness in relation to vocal training in 55 participants, including 27 professionally trained opera singers and 28 non-singers. Results indicated trend-level differences with thicker callosal regions in singers, particularly at the anterior-posterior midbody border and the isthmus, though these did not survive corrections for multiple comparisons. However, a significant negative correlation between age at first singing lesson and callosal thickness was found in the anterior third (rostrum, genu, rostral body), the anterior-posterior midbody border, and the isthmus, suggesting that early vocal training facilitates lasting neuroplastic adaptations in these regions. Additionally, a positive trend between years of professional singing and greater thickness in the midbody was observed but did not remain significant after correction for multiple comparisons. Greater callosal thickness likely enhances interhemispheric connectivity to meet the demands of operatic performance, highlighting the adaptability of the corpus callosum to early, sustained sensorimotor training. By extending evidence from instrumental musicians to singers, these results underscore developmental timing as a key factor in how sensorimotor training shapes brain structure.

## 1. Introduction

Professional musicians are well-known for their astounding levels of fine-motor skills, virtuosically displayed both visually and acoustically as they excel in playing their instrument. Commencing formal education in early childhood, they dedicate years of practice to developing, perfecting, and maintaining the execution of highly complex motor sequences with temporal accuracies within the range of just a few milliseconds (Jabusch et al. 2009; Braun Janzen et al. 2014; Penhune 2022). Linked to these extraordinary levels of sensorimotor and auditory skill, research involving musicians has contributed profound insights into the brain’s remarkable capacity for adaptation (Jancke 2009; Herholz and Zatorre 2012; Leipold et al. 2021). The correspondence between changes in structure, function, and behavior not only underscores the brain’s role in subserving the cognitive and motor demands required for performing music but also highlights the unique ways in which specialized occupational training can shape its architecture (Gärtner et al. 2013; Wu et al. 2020; Olszewska et al. 2021).

Despite the substantial research on use-dependent neural adaptations in instrumental musicians (Munte et al. 2002; Schneider et al. 2002; Gaser and Schlaug 2003; Bermudez and Zatorre 2005; Jancke 2009; James et al. 2014; Groussard et al. 2014; Sato et al. 2015; Elmer et al. 2016; Karpati et al. 2017; Olszewska et al. 2021; Leipold et al. 2021; Penhune 2022), neural plasticity in response to vocal training has been understudied (Halwani et al. 2011; Kleber et al. 2016; Wang et al. 2019; Perron et al. 2021). Singers, unlike instrumentalists, cannot rely on visual guidance to refine their motor skills; instead, they must coordinate complex respiratory, laryngeal, and articulatory motor systems, which undergo significant maturation over an extended period of speech motor development (Smith and Zelaznik 2004; Sadagopan and Smith 2008; Ross et al. 2011). Moreover, singing training demands an elevated level of temporal and tonal motor precision within a musical framework, which significantly exceeds that required for speech (Zatorre and Baum 2012), placing unique demands on the neural circuits involved in sensorimotor integration (Kleber et al. 2013, 2017). Singing experience thus offers a unique window into use-dependent plasticity within brain regions frequently associated with learning-dependent neural plasticity in musicians, such as the corpus callosum (Schlaug et al. 1995; Lee et al. 2003; Steele et al. 2013; Leipold et al. 2021). Given its critical role in interhemispheric communication (Roland et al. 2017), extensive training with a musical instrument during early developmental periods has been associated with enduring changes in the callosal white-matter microstructure and connectivity (Penhune 2022). Understanding such adaptations in singers may offer new insights into the neural underpinnings of sensorimotor control in the speech-motor system (Jurgens 2002; Simonyan and Horwitz 2011), with broader implications for specialized sensorimotor training on callosal structure.

Considering the unique sensorimotor demands of singing (Harris et al. 2023) and the corpus callosum’s essential function in interhemispheric communication within the speech motor system (Friederici et al. 2007; Kort et al. 2014, 2016; Belyk et al. 2015; Neef et al. 2015), this study investigates the structural characteristics of the corpus callosum among professionally trained opera singers compared to non-singers. We hypothesize that the complex vocal motor training required to match style-specific sound targets within a musical framework while concurrently maintaining speech intelligibility, as required in operatic singing, will result in greater callosal thickness in trained singers within callosal subregions associated with interhemispheric sensorimotor communication. Given that the corpus callosum matures during childhood and adolescence (van der Knaap and van der Ham 2011), we further posit that such greater thickness can particularly be observed when individuals began formal singing training early in life.

## 2. Methods

### 2.1 Study Sample

We enlisted 55 right-handed individuals with no reported history of neurological or psychiatric disorders. Handedness was determined using the Edinburgh Handedness Inventory (Oldfield 1971). The participants were classified into two distinct cohorts: a group of professional classical singers (n = 27; mean age = 26.6 years; age range = 20–34 years, including 19 females) and a control group without formal vocal training (n = 28; mean age = 24.9 years; age range = 21–31 years; 21 females).

The singer cohort included members from prominent local musical institutions: two from the Stuttgart State Opera, one from the SWR Radio Vocal Ensemble Stuttgart, and twenty-four from the State University of Music and Performing Arts Stuttgart. The latter group was composed of students engaged in advanced studies in vocal performance or opera. On average, these singers began their formal training at 15.7 years old (SD = 3.7; range = 7–25 years), accumulating an average of 10.4 years of professional experience (SD = 4.2; range = 3–23 years) by the study’s commencement, and dedicating about 17.1 hours per week to singing practice (SD = 6.6; range = 5– 30 hours).

The control group was recruited from the academic community at the University of Tübingen, primarily from the faculties of psychology and medicine. To maintain comparability with the singer group, controls with modest instrumental training were included, with an average musical practice of 3.4 years (range = 1–8 years). However, controls with any extensive singing involvement—defined as more than 5 hours of singing per week—or any formal vocal training were excluded. Written informed consent was obtained from all subjects prior to participation, in accordance with the Declaration of Helsinki. The institutional ethics review board of the Medical Faculty at the University of Tübingen approved the study protocol.

### 2.2 Data acquisition and Preprocessing

Brain Images were obtained using a 1.5 T Sonata whole-body scanner (Siemens Medical Systems, Erlangen, Germany) with a whole-body coil for transmission and an 8-channel phased-array head coil for reception. We used a T1-weighted 3D magnetization prepared rapid gradient echo (MPRAGE) sequence with an isotropic spatial resolution of 1 mm^3^. For each participant, 176 slices were acquired with an image matrix size of 256 × 256 and a FOV = 256 mm × 256 mm^2^. Other image acquisition parameters were TR / TE / TI = 1300 / 3.19 / 660 ms, bandwidth = 190 Hz/Px, and flip-angle = 15°. Head-motion was minimized during scanning with a rubber-foam restraint.

All brain images were pre-processed in SPM12 (http://www.fil.ion.ucl.ac.uk/spm) and the CAT12 toolbox (Gaser et al. 2024) applying corrections for magnetic field inhomogeneities and spatial alignment using rigid-body transformations. In addition, the total intracranial volume (TIV) was estimated for each subject by classifying images as gray matter (GM), white matter (WM), and cerebrospinal fluid (CSF) and adding the sub-volumes of these compartments (TIV = GM + WM + CSF).

### 2.3 Callosal Thickness Estimation

Using the preprocessed images, the corpus callosum was manually outlined by one rater (C.D.) in each brain’s midsagittal section (Luders et al. 2007a). The callosal traces were extracted and automatically processed in a number of successive steps (Luders et al. 2011, 2014, 2018). More specifically, the callosal outlines were separated into 100 nodes and re-sampled at regular intervals rendering the discrete points comprising the two boundaries spatially uniform. Then, a new midline curve was created by calculating the 2D average from the 100 equidistant nodes representing the upper and the lower callosal boundaries. Finally, the distances between the 100 nodes of the upper as well as the lower callosal boundaries to the 100 nodes of the midline curve were calculated.

### 2.4 Statistical Analyses

The statistical analyses were conducted in Matlab (The MathWorks, Natick, MA) using a mass-univariate general linear model. The calculated point-wise callosal distances constituted the dependent variable, group the independent variable, and age as well as TIV the variables of no interest. In addition to the group comparison (opera singers vs. controls) we conducted two correlation analyses within the opera singers, examining (I) the link between callosal thickness and the age at the first singing lesson, and (II) the link between callosal thickness and the years of professional singing experience. Again, age and TIV were considered variables of no interest. To control for multiple comparisons, a Monte Carlo simulation using 10,000 permutations was employed for all analyses, as previously established (Thompson et al. 2004; Luders et al. 2009; Anastasopoulou et al. 2017).

## 3. Results

Opera singers had significantly thicker corpora callosa than controls at the border between anterior and posterior midbody as well as at the isthmus, but none of these effects survived corrections for multiple comparisons. In contrast, there was no region where controls had significantly thicker corpora callosa than professional singers, even at uncorrected significance levels.

With respect to the age at the first singing lesson, we revealed significant negative correlations (i.e., the younger the age, the thicker the corpus callosum). Effects survived corrections for multiple comparisons and were evident within the anterior third (rostrum, genu, rostral body), at the border between anterior and posterior midbody, as well as at the isthmus.

With respect to the years of the professional singing experience, we revealed significant positive correlations (i.e., the more years, the thicker the corpus callosum). This effect was evident at the border between anterior and posterior midbody but did not survive corrections for multiple comparisons.

**Fig 1.**
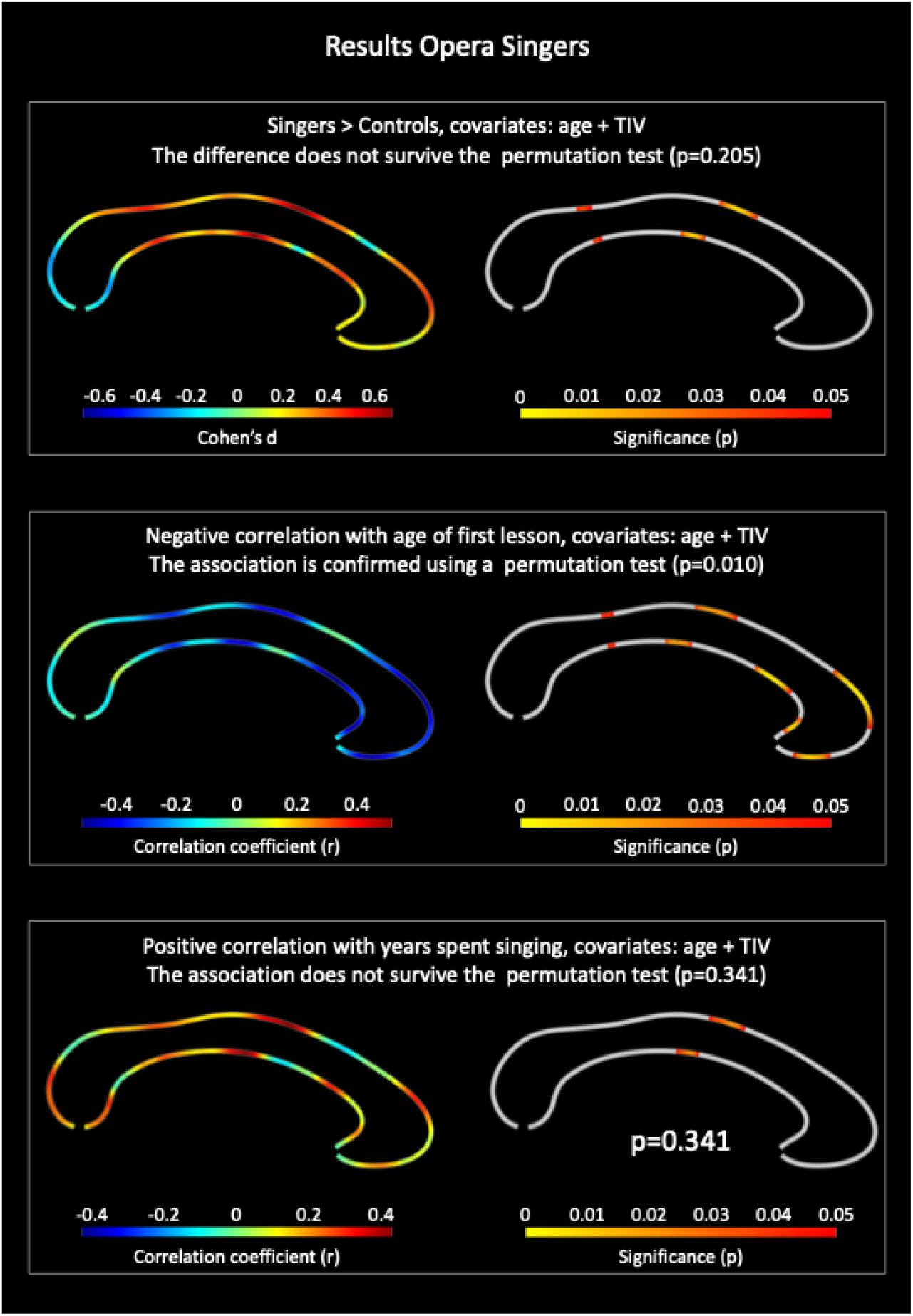
Study Findings. Upper panel – group differences: Thicker corpora callosal in opera singers than controls. Middle panel – negative correlations: The younger the age at the first singing lesson, the thicker the corpus callosum. Bottom panel – positive correlations: The more years of singing experience, the thicker the corpus callosum. The color bar encodes significance (p) at uncorrected levels. The outcomes pertaining to the negative correlation (middle panel) survived corrections for multiple comparisons at p=0.01

## 4. Discussion

This study reveals significant neuroplastic adaptations within the corpus callosum for early-trained classical singers, indicating that an earlier start to vocal training is associated with greater callosal thickness in regions essential for executive functions, motor coordination, auditory processing, and sensory integration. Additionally, years of professional singing experience showed a trend toward increased thickness in the midbody region, though this finding did not reach statistical significance. These observations suggest enhanced interhemispheric connectivity in early-trained singers, likely reflecting the higher sensorimotor demands of singing compared to speech production. Together, our findings underscore the crucial role of developmental timing in maximizing the neuroplastic potential of the corpus callosum through professional vocal training, extending evidence of use-dependent plasticity from instrumental musicians to vocal artists.

### Functional Organization of the Corpus Callosum

The corpus callosum is the largest white matter structure in the brain and facilitates interhemispheric communication by connecting various cortical areas across the left and right cerebral hemispheres mostly in a homotopic fashion (Luders et al. 2018). As such, it has been subdivided into functionally distinct subregions: the *rostrum* connects the fronto-basal cortex, supporting executive functions and emotional regulation; the *genu* links the prefrontal and anterior cingulate cortices, involved in decision-making, attention, and social behavior; the *rostral body* connects the prefrontal cortex and higher-order premotor areas, including parts of the insula; the *anterior midbody* links supplementary motor and premotor areas essential for planning and coordinating complex movements; the *posterior midbody* associates with premotor and partially with primary motor areas; the *isthmus* connects primary motor, sensory, and auditory regions; and finally, the *splenium*, connecting the parietal, temporal, and occipital lobes, plays a key role in integrating sensory information across hemispheres (Witelson 1989; Zarei et al. 2006; Hofer and Frahm 2006; van der Knaap and van der Ham 2011). Their role in the regulation of communication between hemispheres is influenced by callosal fiber composition (size and density), which determines interhemispheric transfer times (Aboitiz et al. 1992a, b; Clarke and Zaidel 1994). The thickness of the corpus callosum (CC) moreover contributes to interhemispheric integration. Thicker callosal regions, particularly in posterior and anterior sections, are associated with greater hemispheric connectivity, facilitating more efficient transfer of information across hemispheres (Luders et al. 2007b, 2012). Notably, the corpus callosum develops and matures throughout childhood and adolescence with increasing myelination, reflecting corresponding changes in cognitive and motor functions (van der Knaap and van der Ham 2011).

### Neuroplastic Adaptations in the Corpus Callosum of Instrumental Musicians

Early magnetic resonance imaging (MRI) studies reported a larger anterior corpus callosum region in professional instrumentalists, particularly those who began training before age 7, suggesting that sensorimotor training during sensitive periods supports callosal maturation (Schlaug et al. 1995; Ozturk et al. 2002; Lee et al. 2003). Subsequent diffusion-weighted imaging (DWI) studies further revealed that early-trained musicians typically exhibit greater interhemispheric connectivity in the isthmus and splenium, followed by the anterior and posterior midbody, with fewer findings in the rostral body and genu (Schmithorst and Wilke 2002; Bengtsson et al. 2005; Imfeld et al. 2009; Steele et al. 2013; Leipold et al. 2021). Moreover, current training intensity in middle-aged musicians appears to influence the isthmus and splenium, linking enhanced interhemispheric communication to improved bimanual fine-motor control (Gärtner et al. 2013). Collectively, these studies underscore the critical role of the corpus callosum in coordinating bilateral brain functions necessary for auditory processing and sensorimotor integration in highly trained instrumentalists (Leipold et al. 2021; Penhune 2022).

### Neuroplastic Adaptations in the Corpus Callosum of Classical Singers

The specialized motor skills of instrumentalists differ notably from those of singers, who fine-tune their speech motor system to meet the heightened precision demands of music production. Unlike limb motor control, which is strongly lateralized, coordination of the tongue, pharynx, and larynx in singers is bilaterally controlled (Jurgens 2002; Simonyan and Horwitz 2011; Pilurzi et al. 2013). Moreover, the bilateral human larynx motor cortex (LMC), essential for precise pitch control (Dichter et al. 2018)—makes direct monosynaptic connections to the brainstem’s nucleus ambiguous, which causes muscles involved in vocal production to contract. These descending fibers from each LMC cross incompletely at the medulla oblongata, allowing one hemisphere to engage both left and right vocal folds, with a contralateral predominance (Jurgens 2002; Pilurzi et al. 2013; Simonyan 2014; Simonyan et al. 2016).

While this anatomical setup suggests that input from one hemisphere would suffice to control phonation, interhemispheric communication still plays a crucial role in coordinating the motor production and perception processes underlying speech (Friederici et al. 2007; Belyk et al. 2015; Neef et al. 2015). Specifically, integrating auditory feedback with motor control primarily involves the left hemisphere for executing and planning speech motor functions, while the right motor cortex contributes to modulation and fine-tuning (Kort et al. 2014, 2016). Additionally, the left LMC is predominantly active during speech perception, while the right LMC shows greater engagement in cognitively demanding tasks (Liang et al. 2023), suggesting differential processing of non-linguistic musical voice stimuli (Leveque et al. 2013). This aligns with the observation that singing, in contrast to speaking, involves more extensive bilateral brain activation, with a notable increase in right hemisphere contribution (Callan et al. 2006; Ozdemir et al. 2006). This heightened interhemispheric coordination likely supports the integration of auditory and motor processes needed for vocal production, which must adhere to specific temporal and timbral constraints within a musical framework (Zatorre and Baum 2012; Zatorre 2013; Albouy et al. 2020). Previous studies by Kleber et al. (2010, 2013, 2016, 2017) demonstrate enhanced integration of auditory, somatosensory, and motor information in expert singers, aligning with evidence of structural and functional adaptations within the brain. These studies largely focus on the specific integration necessary for the fine motor control, auditory precision, and feedback processing required for singing. The current findings of a thicker corpus callosum in the midbody and isthmus align with these reports, suggesting increased capacity for interhemispheric communication in opera singers, particularly among those who began training at a young age. Additionally, recent findings indicate that singing training in individuals with aphasia can induce structural neuroplasticity in the corpus callosum, potentially supporting enhanced interhemispheric communication essential for coordinated vocal production (Sihvonen et al. 2024)

The negative association between callosal thickness and the age at first vocal lesson in the rostrum, genu, and anterior midbody suggests that early training may enhance interhemispheric connectivity in regions involved in executive function, attention, and motor planning. This association, particularly evident in the anterior midbody, aligns with previous findings of stronger engagement in brain areas associated with higher-level attention, performance monitoring, and motor planning in singers, including the dorsolateral prefrontal cortex, supplementary motor area (SMA), and key regions of the salience network (Kleber et al. 2007, 2010, 2017). Enhanced interhemispheric communication via these sub-regions may thus support the complex sensorimotor planning and top-down control processes to facilitate precise tonal, temporal, and timbral sound production (Sundberg et al. 1993; Sundberg 1994; Dromey et al. 2015; Scherer et al. 2017; Dichter et al. 2018).

### Comparative Insights between Singers and Instrumentalists

Despite the bilateral cortical representation of phonation-related muscles in singers, our findings reveal both shared and unique patterns of use-dependent plasticity in the corpus callosum, similar to adaptations seen in instrumentalists. Greater callosal thickness in the isthmus suggests a domain-general role in auditory and motor integration essential for both vocal and instrumental music production. Additionally, findings at the border between the anterior and posterior midbody indicate enhanced integration of somatosensory and motor information, paralleling adaptations seen in instrumentalists that support fine motor control and bimanual coordination. This pattern aligns with prior studies showing that expert singers exhibit task- and experience-specific activation patterns in motor and sensory regions (Kleber et al. 2010, 2013, 2016, 2017). However, the increased connectivity observed in singers within the rostrum, genu, and rostral body—regions linked to prefrontal processing, emotional regulation, and executive functions—differs from typical findings in instrumentalists (Schlaug et al. 1995; Lee et al. 2003; Steele et al. 2013; Leipold et al. 2021), suggesting that they may reflect the unique demands of vocal control and expressive performance in singers.

Interestingly, patterns of interhemispheric connectivity and inhibition also vary across types of instrumentalists, pointing to additional experience-dependent effects. Structural enlargements in the corpus callosum are generally associated with reduced interhemispheric inhibition and enhanced processing (Ridding et al. 2000). However, findings reveal stronger left-to-right interhemispheric inhibition (IHI) in string players compared to pianists, whose IHI is comparable to that of non-musicians (Vollmann et al. 2014). In singers, interhemispheric inhibitory motor interactions are more pronounced during singing than in speech or humming tasks (Lo and Fook-Chong 2004), paralleling the refined bilateral control essential for vocal performance. Additionally, studies on bilingual proficiency suggest that complex vocal tasks may enhance interhemispheric connectivity due to increased cognitive demands, as seen in the strengthened connectivity between Brodmann Areas 44 and 45 (Sander et al. 2023). Together, these findings indicate specific experience-dependent effects on the corpus callosum, where enhanced connectivity in the rostrum, genu, and rostral body likely supports advanced cognitive control and motor planning necessary for adjusting vocal behaviors.

## Conclusions

This study demonstrates the dynamic neuroplasticity of the corpus callosum in response to specialized vocal training, with notable findings on how the age of training onset influences callosal development. Increased callosal thickness in regions such as the rostrum, genu, rostral body, anterior-posterior midbody, and isthmus suggests that early training enhances interhemispheric connectivity essential for the complex sensorimotor and cognitive demands of artistic vocal performance.

These structural adaptations likely support the precise sensorimotor, auditory, and cognitive-emotional integration required in operatic singing, while also revealing unique aspects of callosal development in singers compared to instrumentalists. The findings underscore the corpus callosum’s central role in coordinating high-level motor and cognitive processes and highlight the impact of vocal training on the development of these interhemispheric pathways. Differences in callosal adaptations between singers and instrumentalists emphasize the specificity of training demands, offering insights into how varied forms of musical expertise uniquely shape brain structure over time.

## Statements & Declarations

## Author Contributions

BK and ML contributed to the study conception and design, and all authors made substantial contributions to the analysis or interpretation of data. Material preparation and data acquisition were performed by BK and ML. Data analyses were performed by FK, EL, and CD. The first draft of the manuscript was written by BK, AZ, EL, and FK, and all authors contributed to drafting the work or revising it critically by commenting on previous versions of the manuscript. All authors read, reviewed, and approved the final manuscript.

## Data Availability

The datasets generated during and/or analyzed during the current study are not publicly available due to European GDPR restrictions but are available from the corresponding author on reasonable request.

## Acknowledgments

We thank the Staatliche Hochschule für Musik und Darstellende Kunst Stuttgart and the Staatsoper Stuttgart (Germany) for their support.

## References

Aboitiz F, Scheibel AB, Fisher RS, Zaidel E (1992a) Fiber composition of the human corpus callosum. Brain Res 598:143– 153. 10.1016/0006-8993(92)90178-C

Aboitiz F, Scheibel AB, Fisher RS, Zaidel E (1992b) Individual differences in brain asymmetries and fiber composition in the human corpus callosum. Brain Res 598:154–161. 10.1016/0006-8993(92)90179-D

Albouy P, Benjamin L, Morillon B, Zatorre RJ (2020) Distinct sensitivity to spectrotemporal modulation supports brain asymmetry for speech and melody. Science 367:1043–1047. 10.1126/science.aaz3468

Anastasopoulou S, Kurth F, Luders E, Savic I (2017) Generalized epilepsy syndromes and callosal thickness: Differential effects between patients with juvenile myoclonic epilepsy and those with generalized tonic-clonic seizures alone. Epilepsy Res 129:74–78. 10.1016/j.eplepsyres.2016.11.008

Belyk M, Kraft SJ, Brown S (2015) Stuttering as a trait or state – an ALE meta-analysis of neuroimaging studies. Eur J Neurosci 41:275–284. 10.1111/ejn.12765

Bengtsson SL, Nagy Z, Skare S, et al (2005) Extensive piano practicing has regionally specific effects on white matter development. Nat Neurosci 8:1148–50. 10.1038/nn1516

Bermudez P, Zatorre RJ (2005) Differences in gray matter between musicians and nonmusicians. Ann N Y Acad Sci 1060:395–9. 10.1196/annals.1360.057

Braun Janzen T, Thompson WF, Ranvaud R (2014) A developmental study of the effect of music training on timed movements. Front Hum Neurosci 8:801. 10.3389/fnhum.2014.00801

Callan DE, Tsytsarev V, Hanakawa T, et al (2006) Song and speech: brain regions involved with perception and covert production. NeuroImage 31:1327–42. 10.1016/j.neuroimage.2006.01.036

Clarke JM, Zaidel E (1994) Anatomical-behavioral relationships: corpus callosum morphometry and hemispheric specialization. Behav Brain Res 64:185–202. 10.1016/0166-4328(94)90131-7

Dichter BK, Breshears JD, Leonard MK, Chang EF (2018) The Control of Vocal Pitch in Human Laryngeal Motor Cortex. Cell 174:21–31 e9. 10.1016/j.cell.2018.05.016

Dromey C, Holmes SO, Hopkin JA, Tanner K (2015) The Effects of Emotional Expression on Vibrato. J Voice 29:170–181. 10.1016/j.jvoice.2014.06.007

Elmer S, Hanggi J, Jancke L (2016) Interhemispheric transcallosal connectivity between the left and right planum temporale predicts musicianship, performance in temporal speech processing, and functional specialization. Brain Struct Funct 221:331–44. 10.1007/s00429-014-0910-x

Friederici AD, von Cramon DY, Kotz SA (2007) Role of the Corpus Callosum in Speech Comprehension: Interfacing Syntax and Prosody. Neuron 53:135–145. 10.1016/j.neuron.2006.11.020

Gärtner H, Minnerop M, Pieperhoff P, et al (2013) Brain morphometry shows effects of long-term musical practice in middle-aged keyboard players. Front Psychol 4:

Gaser C, Dahnke R, Thompson PM, et al (2024) CAT: a computational anatomy toolbox for the analysis of structural MRI data. GigaScience 13:giae049. 10.1093/gigascience/giae049

Gaser C, Schlaug G (2003) Brain structures differ between musicians and non-musicians. J Neurosci 23:9240–9245

Groussard M, Viader F, Landeau B, et al (2014) The effects of musical practice on structural plasticity: the dynamics of grey matter changes. Brain Cogn 90:174–80. 10.1016/j.bandc.2014.06.013

Halwani GF, Loui P, Ruber T, Schlaug G (2011) Effects of practice and experience on the arcuate fasciculus: comparing singers, instrumentalists, and non-musicians. Front Psychol 2:156. 10.3389/fpsyg.2011.00156

Harris I, Niven EC, Griffin A, Scott SK (2023) Is song processing distinct and special in the auditory cortex? Nat Rev Neurosci 24:711–722. 10.1038/s41583-023-00743-4

Herholz SC, Zatorre RJ (2012) Musical training as a framework for brain plasticity: behavior, function, and structure. Neuron 76:486–502. 10.1016/j.neuron.2012.10.011

Hofer S, Frahm J (2006) Topography of the human corpus callosum revisited—Comprehensive fiber tractography using diffusion tensor magnetic resonance imaging. NeuroImage 32:989–994. 10.1016/j.neuroimage.2006.05.044

Imfeld A, Oechslin MS, Meyer M, et al (2009) White matter plasticity in the corticospinal tract of musicians: A diffusion tensor imaging study. NeuroImage 46:600–607. 10.1016/j.neuroimage.2009.02.025

Jabusch HC, Alpers H, Kopiez R, et al (2009) The influence of practice on the development of motor skills in pianists: a longitudinal study in a selected motor task. Hum Mov Sci 28:74–84. 10.1016/j.humov.2008.08.001

James CE, Oechslin MS, Van De Ville D, et al (2014) Musical training intensity yields opposite effects on grey matter density in cognitive versus sensorimotor networks. Brain Struct Funct 219:353–366. 10.1007/s00429-013-0504-z

Jancke L (2009) The plastic human brain. Restor Neurol Neurosci 27:521–38. 10.3233/RNN-2009-0519

Jurgens U (2002) Neural pathways underlying vocal control. Neurosci Biobehav Rev 26:235–58

Karpati FJ, Giacosa C, Foster NEV, et al (2017) Dance and music share gray matter structural correlates. Brain Res 1657:62– 73. 10.1016/j.brainres.2016.11.029

Kleber B, Birbaumer N, Veit R, et al (2007) Overt and imagined singing of an Italian aria. NeuroImage 36:889–900. 10.1016/j.neuroimage.2007.02.053

Kleber B, Friberg A, Zeitouni A, Zatorre R (2017) Experience-dependent modulation of right anterior insula and sensorimotor regions as a function of noise-masked auditory feedback in singers and nonsingers. NeuroImage 147:97–110. 10.1016/j.neuroimage.2016.11.059

Kleber B, Veit R, Birbaumer N, et al (2010) The brain of opera singers: experience-dependent changes in functional activation. Cereb Cortex 20:1144–52. 10.1093/cercor/bhp177

Kleber B, Veit R, Moll CV, et al (2016) Voxel-based morphometry in opera singers: Increased gray-matter volume in right somatosensory and auditory cortices. NeuroImage 133:477–483. 10.1016/j.neuroimage.2016.03.045

Kleber B, Zeitouni AG, Friberg A, Zatorre RJ (2013) Experience-dependent modulation of feedback integration during singing: role of the right anterior insula. J Neurosci 33:6070–80. 10.1523/JNEUROSCI.4418-12.2013

Kort NS, Cuesta P, Houde JF, Nagarajan SS (2016) Bihemispheric network dynamics coordinating vocal feedback control. Hum Brain Mapp 37:1474–85. 10.1002/hbm.23114

Kort NS, Nagarajan SS, Houde JF (2014) A bilateral cortical network responds to pitch perturbations in speech feedback. NeuroImage 86:525–35. 10.1016/j.neuroimage.2013.09.042

Lee DJ, Chen Y, Schlaug G (2003) Corpus callosum: musician and gender effects. NeuroReport 14:205–9. 10.1097/01.wnr.0000053761.76853.41

Leipold S, Klein C, Jäncke L (2021) Musical Expertise Shapes Functional and Structural Brain Networks Independent of Absolute Pitch Ability. J Neurosci Off J Soc Neurosci 41:2496–2511. 10.1523/JNEUROSCI.1985-20.2020

Leveque Y, Muggleton N, Stewart L, Schon D (2013) Involvement of the larynx motor area in singing-voice perception: a TMS study(dagger). Front Psychol 4:418. 10.3389/fpsyg.2013.00418

Liang B, Li Y, Zhao W, Du Y (2023) Bilateral human laryngeal motor cortex in perceptual decision of lexical tone and voicing of consonant. Nat Commun 14:4710. 10.1038/s41467-023-40445-0

Lo YL, Fook-Chong S (2004) Ipsilateral and contralateral motor inhibitory control in musical and vocalization tasks. Exp Brain Res 159:258–262. 10.1007/s00221-004-2032-9

Luders E, Di Paola M, Tomaiuolo F, et al (2007a) Callosal morphology in Williams syndrome: a new evaluation of shape and thickness. Neuroreport 18:203–207. 10.1097/WNR.0b013e3280115942

Luders E, Narr KL, Bilder RM, et al (2007b) Positive correlations between corpus callosum thickness and intelligence. NeuroImage 37:1457–1464. 10.1016/j.neuroimage.2007.06.028

Luders E, Narr KL, Hamilton LS, et al (2009) Decreased callosal thickness in attention-deficit/hyperactivity disorder. Biol Psychiatry 65:84–88. 10.1016/j.biopsych.2008.08.027

Luders E, Phillips OR, Clark K, et al (2012) Bridging the hemispheres in meditation: Thicker callosal regions and enhanced fractional anisotropy (FA) in long-term practitioners. NeuroImage 61:181–187. 10.1016/j.neuroimage.2012.02.026

Luders E, Thompson PM, Kurth F (2018) Morphometry of the Corpus Callosum. In: Spalletta G, Piras F, Gili T (eds) Brain Morphometry. Springer, New York, NY, pp 131–142

Luders E, Thompson PM, Narr KL, et al (2011) The link between callosal thickness and intelligence in healthy children and adolescents. NeuroImage 54:1823–1830. 10.1016/j.neuroimage.2010.09.083

Luders E, Toga AW, Thompson PM (2014) Why size matters: Differences in brain volume account for apparent sex differences in callosal anatomy: The sexual dimorphism of the corpus callosum. NeuroImage 84:820–824. 10.1016/j.neuroimage.2013.09.040

Munte TF, Altenmüller E, Jancke L (2002) The musician’s brain as a model of neuroplasticity. Nat Rev Neurosci 3:473–8. 10.1038/nrn843

Neef NE, Anwander A, Friederici AD (2015) The Neurobiological Grounding of Persistent Stuttering: from Structure to Function. Curr Neurol Neurosci Rep 15:63. 10.1007/s11910-015-0579-4

Oldfield RC (1971) The assessment and analysis of handedness: the Edinburgh inventory. Neuropsychologia 9:97–113

Olszewska AM, Gaca M, Herman AM, et al (2021) How Musical Training Shapes the Adult Brain: Predispositions and Neuroplasticity. Front Neurosci 15:

Ozdemir E, Norton A, Schlaug G (2006) Shared and distinct neural correlates of singing and speaking. NeuroImage 33:628– 35. 10.1016/j.neuroimage.2006.07.013

Ozturk AH, Tascioglu B, Aktekin M, et al (2002) Morphometric comparison of the human corpus callosum in professional musicians and non-musicians by using in vivo magnetic resonance imaging. J Neuroradiol 29:29–34

Penhune VB (2022) Understanding Sensitive Period Effects in Musical Training. In: Andersen SL (ed) Sensitive Periods of Brain Development and Preventive Interventions. Springer International Publishing, Cham, pp 167–188

Perron M, Theaud G, Descoteaux M, Tremblay P (2021) The frontotemporal organization of the arcuate fasciculus and its relationship with speech perception in young and older amateur singers and non-singers. Hum Brain Mapp 42:3058– 3076. 10.1002/hbm.25416

Pilurzi G, Hasan A, Saifee TA, et al (2013) Intracortical circuits, sensorimotor integration and plasticity in human motor cortical projections to muscles of the lower face. J Physiol 591:1889–1906. 10.1113/jphysiol.2012.245746

Ridding MC, Brouwer B, Nordstrom MA (2000) Reduced interhemispheric inhibition in musicians. Exp Brain Res 133:249– 253. 10.1007/s002210000428

Roland JL, Snyder AZ, Hacker CD, et al (2017) On the role of the corpus callosum in interhemispheric functional connectivity in humans. Proc Natl Acad Sci U S A 114:13278–13283. 10.1073/pnas.1707050114

Ross LA, Molholm S, Blanco D, et al (2011) The development of multisensory speech perception continues into the late childhood years. Eur J Neurosci 33:2329–2337. 10.1111/j.1460-9568.2011.07685.x

Sadagopan N, Smith A (2008) Developmental changes in the effects of utterance length and complexity on speech movement variability. J Speech Lang Hear Res JSLHR 51:1138–1151. 10.1044/1092-4388(2008/06-0222)

Sander K, Chai X, Barbeau EB, et al (2023) Interhemispheric functional brain connectivity predicts new language learning success in adults. Cereb Cortex 33:1217–1229. 10.1093/cercor/bhac131

Sato K, Kirino E, Tanaka S (2015) A Voxel-Based Morphometry Study of the Brain of University Students Majoring in Music and Nonmusic Disciplines. Behav Neurol 2015:274919. 10.1155/2015/274919

Scherer KR, Sundberg J, Fantini B, et al (2017) The expression of emotion in the singing voice: Acoustic patterns in vocal performance. J Acoust Soc Am 142:1805. 10.1121/1.5002886

Schlaug G, Jäncke L, Huang Y, et al (1995) Increased corpus callosum size in musicians. Neuropsychologia 33:1047–1055. 10.1016/0028-3932(95)00045-5

Schmithorst VJ, Wilke M (2002) Differences in white matter architecture between musicians and non-musicians: a diffusion tensor imaging study. Neurosci Lett 321:57–60. https://doi.org/PiiS0304-3940(02)00054-X Doi 10.1016/S0304-3940(02)00054-X

Schneider P, Scherg M, Dosch HG, et al (2002) Morphology of Heschl’s gyrus reflects enhanced activation in the auditory cortex of musicians. Nat Neurosci 5:688–94. 10.1038/nn871

Sihvonen AJ, Pitkäniemi A, Siponkoski S-T, et al (2024) Structural Neuroplasticity Effects of Singing in Chronic Aphasia. eNeuro 11:. 10.1523/ENEURO.0408-23.2024

Simonyan K (2014) The laryngeal motor cortex: its organization and connectivity. Curr Opin Neurobiol 28:15–21. 10.1016/j.conb.2014.05.006

Simonyan K, Ackermann H, Chang EF, Greenlee JD (2016) New Developments in Understanding the Complexity of Human Speech Production. J Neurosci 36:11440–11448. 10.1523/JNEUROSCI.2424-16.2016

Simonyan K, Horwitz B (2011) Laryngeal motor cortex and control of speech in humans. Neuroscientist 17:197–208. 10.1177/1073858410386727

Smith A, Zelaznik HN (2004) Development of functional synergies for speech motor coordination in childhood and adolescence. Dev Psychobiol 45:22–33. 10.1002/dev.20009

Steele CJ, Bailey JA, Zatorre RJ, Penhune VB (2013) Early musical training and white-matter plasticity in the corpus callosum: evidence for a sensitive period. J Neurosci 33:1282–90. 10.1523/JNEUROSCI.3578-12.2013

Sundberg J (1994) Perceptual aspects of singing. J Voice 8:106–22

Sundberg J, Gramming P, Lovetri J (1993) Comparisons of pharynx, source, formant, and pressure characteristics in operatic and musical theatre singing. J Voice 7:301–10

Thompson PM, Hayashi KM, Sowell ER, et al (2004) Mapping cortical change in Alzheimer’s disease, brain development, and schizophrenia. NeuroImage 23 Suppl 1:S2–18. 10.1016/j.neuroimage.2004.07.071

van der Knaap LJ, van der Ham IJM (2011) How does the corpus callosum mediate interhemispheric transfer? A review. Behav Brain Res 223:211–221. 10.1016/j.bbr.2011.04.018

Vollmann H, Ragert P, Conde V, et al (2014) Instrument specific use-dependent plasticity shapes the anatomical properties of the corpus callosum: a comparison between musicians and non-musicians. Front Behav Neurosci 8:

Wang W, Wei L, Chen N, et al (2019) Decreased Gray-Matter Volume in Insular Cortex as a Correlate of Singers’ Enhanced Sensorimotor Control of Vocal Production. Front Neurosci 13:

Witelson SF (1989) Hand and sex differences in the isthmus and genu of the human corpus callosum. A postmortem morphological study. Brain 112:799–835. 10.1093/brain/112.3.799

Wu H, Yan H, Yang Y, et al (2020) Occupational Neuroplasticity in the Human Brain: A Critical Review and Meta-Analysis of Neuroimaging Studies. Front Hum Neurosci 14:

Zarei M, Johansen-Berg H, Smith S, et al (2006) Functional anatomy of interhemispheric cortical connections in the human brain. J Anat 209:311–320. 10.1111/j.1469-7580.2006.00615.x

Zatorre RJ (2013) Predispositions and plasticity in music and speech learning: neural correlates and implications. Science 342:585–9. 10.1126/science.1238414

Zatorre RJ, Baum SR (2012) Musical melody and speech intonation: singing a different tune. PLoS Biol 10:e1001372. 10.1371/journal.pbio.1001372

